# GaugeFixer: overcoming parameter non-identifiability in models of sequence-function relationships

**DOI:** 10.64898/2025.12.08.693054

**Authors:** Carlos Martí-Gómez, David M. McCandlish, Justin B. Kinney

## Abstract

**Background:** Mathematical models of sequence-function relationships are widely used in computational biology. A key challenge when interpreting these models is that many different choices for model parameters can encode the same sequence-function relationship. These ambiguities, called “gauge freedoms,” must be removed before parameter values can be meaningfully interpreted. Doing this requires imposing additional mathematical constraints on parameter values, a process called “fixing the gauge.” We recently developed mathematical methods for fixing the gauge of a large class of commonly used models. The most direct computational implementation of these methods is often impractical, however, because it requires a projection matrix whose size scales quadratically with the number of model parameters.

**Results:** Here we introduce GaugeFixer, a Python package that exploits the mathematical structure of gauge-fixing projections to achieve log-linear scaling in computation time and linear scaling in memory usage. GaugeFixer thus requires orders of magnitude less time and memory than standard matrix multiplication, and can fix the gauge of models having millions of parameters in seconds. To demonstrate its utility, we used GaugeFixer to analyze an empirical fitness landscape for translation initiation in bacteria. This analysis revealed striking similarities (but also fine-scale differences) in ribosome-binding preferences at different positions relative to the start codon, thereby aiding the interpretation of this otherwise unwieldy landscape.

**Conclusions:** GaugeFixer thus fills an important gap in the computational tools available for interpreting quantitative models of sequence-function relationships.

## Background

Computational biology routinely involves models of sequence-function relationships, i.e., models that describe the quantitative relationship between biological sequences (DNA, RNA, or protein) and measurable activities [1]. Such models are frequently used to predict the locations of transcription factor binding sites in promoters and enhancers [2], the locations of splice sites in pre-mRNA transcripts [3], structural contacts between residues in folded proteins [4], and the effects of human genetic variation on protein function [5]. Another rapidly growing application is the analysis of high-throughput mutagenesis data [6–15].

Sequence-function relationships are commonly represented using generalized one-hot models [16]. In these models, each sequence is encoded as a vector of binary features, each feature indicating the presence or absence of a specific subsequence at a specific set of positions. For example, additive features indicate individual characters (nucleotides or amino acids) at individual positions, pairwise features indicate pairs of characters at pairs of positions, and so on. For each sequence feature, there is a corresponding model parameter that quantifies the effect of that feature on a sequence’s activity. Although these models are linear functions of their parameters, they can accommodate genetic interactions of arbitrary order, and the most general of them can represent arbitrarily complex sequence-function landscapes.

A fundamental challenge in interpreting the parameters of these models is that, except in trivial cases, parameter values are not uniquely determined by the sequence-function landscape that the model describes [17]. This structural non-identifiability is a consequence of the model’s mathematical form and is independent of the type and amount of data used to fit it. It occurs because the binary feature vectors representing all possible sequences span only a proper subspace of the vector space in which they are defined. Consequently, a continuum of parameter vectors yields identical model predictions. These parameter vectors differ from one another along what are called “gauge freedoms”—directions in parameter space that are orthogonal to the binary feature vectors of all possible sequences.

The ambiguity that gauge freedoms produce must be removed before the values of model parameters can be interpreted, or before the parameters of two different models can be compared. This is done by imposing additional mathematical constraints that, together with the landscape itself, select a unique parameter vector. This procedure is called “fixing the gauge,” and each choice of constraints defines a distinct gauge. Different gauges provide different mathematical representations of the same underlying sequence-function relationship, and thus correspond to different interpretations of the model’s parameters.

The importance of gauge freedoms was first recognized in theoretical physics and is very well understood in that context [18]. Until recently, however, gauge freedoms in sequence-function relationships had not received focused attention. A scattered body of work described gauge-fixing approaches for specific modeling applications, e.g., in additive models of transcription factor binding specificity [19–21], pairwise models of protein fitness landscapes [22–29], and all-order models in various settings [30–32]. Gauge freedoms also arose implicitly in the literature concerning alternative parameterizations of genetic interactions [33–39]. A unified understanding of gauge freedoms and gauge-fixing approaches, however, was still lacking.

To address this need, we recently developed a mathematical theory of gauge freedoms in generalized one-hot models [16, 17]. In particular, Posfai et al. [16] introduced a family of gauges that includes nearly all of the gauges used in the prior literature, and that can be imposed by multiplying the vector of model parameters by a projection matrix having a simple mathematical form. However, this operation requires storing and manipulating a dense projection matrix whose size scales quadratically with the number of model parameters. This is impractical for models with more than about ten thousand parameters.

Here we introduce GaugeFixer, an open-source Python package that overcomes this computational limitation. By exploiting the mathematical structure of the projection matrices described by Posfai et al. [16], GaugeFixer achieves log-linear scaling in computation time and linear scaling in memory, enabling rapid gauge fixing for models having millions of parameters. To demonstrate its utility, we applied GaugeFixer to a model for translation initiation that has nearly two million parameters [14, 40]. This analysis illustrates how GaugeFixer can reveal both the shared sequence requirements and the fine-scale differences among ribosome-binding sites at different positions relative to the start codon, patterns that are otherwise difficult to discern in a landscape of this complexity.

## Technical Background

### Models

We consider generalized one-hot models that assign a scalar activity *f*(*s*) to sequences *s* of fixed length *L*. Of particular interest among these are all-order models, which describe interactions of every order up through *L*. Such models can represent any possible sequence-function relationship, but the number of parameters they require grows exponentially with sequence length. It is therefore common to use models that include only interactions up to a specified order (*K*-order models) or only interactions between nearby positions (*K*-adjacent models). The number of parameters in these models grows only polynomially with sequence length (as *L*^*K*^ for *K*-order models and linearly in *L* for *K*-adjacent models), making them tractable for much longer sequences. All-order, *K*-order, and *K*-adjacent models are examples of hierarchical models, a subset of generalized one-hot models with a mathematical structure that greatly simplifies the task of fixing the gauge. See Implementation for details.

### Minimal gauge fixing example

We now illustrate the process of gauge fixing using a minimal example of a sequence-function landscape. Consider sequences of length *L* = 1 built from a two-character alphabet {A, B}. There are two possible sequences, *s* = *A* and *s* = *B*, so the landscape has only two degrees of freedom. The corresponding all-order model, however, has three parameters—a constant parameter (*θ*_0_), which applies to all sequences, and two additive parameters (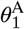 and 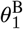) that describe the effects of the two possible characters at position 1 in the sequence. These parameters determine the landscape via:

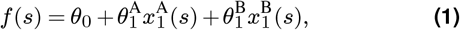

where 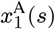 equals 1 if *s* = *A* and equals 0 otherwise; 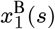 is defined similarly. The model has one more parameter than the landscape has degrees of freedom, and so exhibits one gauge freedom. Fixing the gauge involves imposing an additional constraint that removes this gauge freedom.

Figure 1 illustrates this. Consider the parameter vector 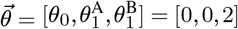. This choice yields the sequence-function landscape

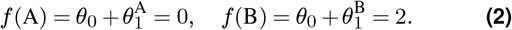

**Fig. 1.**
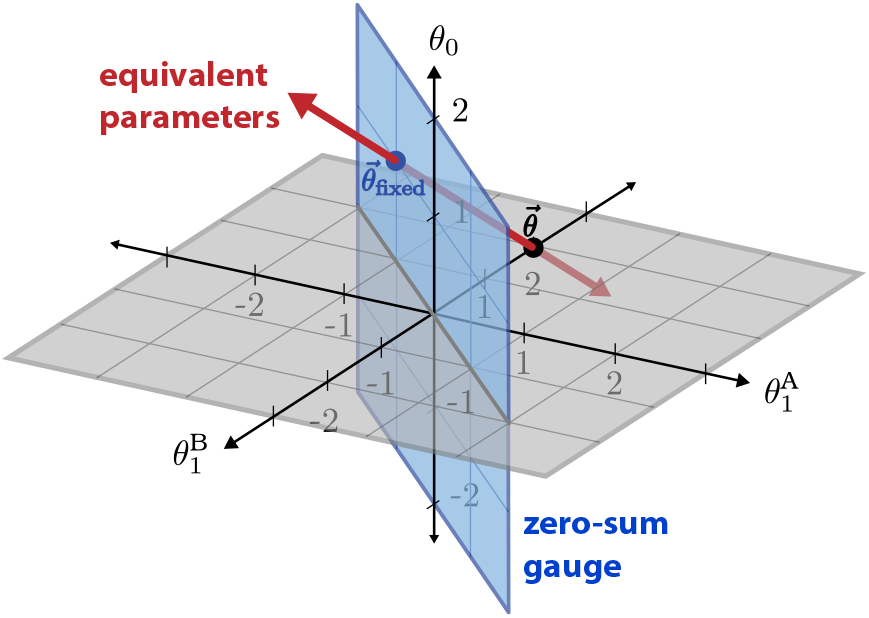
Minimal example of gauge fixing. Shown is the three-dimensional space of parameter vectors 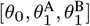, corresponding to an all-order model for sequences of length *L* = 1 built from the two-character alphabet {A, B}. Black dot: the parameter vector 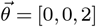. Red line: parameter vectors equivalent to 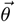. Blue plane: parameter vectors in the zero-sum gauge, i.e., those satisfying 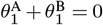. Blue dot: gauge-fixed parameters 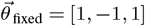.

Any parameter vector of the form [*z*, −*z*,2 − *z*], where z is a real number, will also yield the same landscape. This set of equivalent parameters forms a line within the three-dimensional space of parameter vectors; fixing the gauge amounts to selecting a single point on this line. One way to make this selection is by adopting the zero-sum gauge, which requires that the additive parameters at each position sum to zero (here,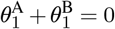).The blue plane in Figure 1 shows the parameter vectors satisfying this constraint. The resulting gauge-fixed parameters, 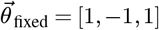, lie at the intersection of the line of equivalent parameters with the zero-sum plane. Importantly, no other parameter vector in the zero-sum gauge yields this same landscape. Adopting this gauge also makes the individual parameter values interpretable: *θ*_0_ = 1 represents the average activity across all possible sequences, whereas 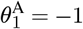 and 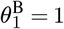 represent the contributions of characters *A* and *B*, respectively, relative to this average value.

### Families of gauges

We previously introduced a family of gauges for all-order models [16]. These gauges are parameterized by two quantities: a non-negative real number λ and a probability distribution *π* over sequences that factorizes across positions. The parameter λ controls how explained variance is distributed across interaction orders, while *π* specifies the sequence distribution over which that variance is computed. This λ, *π* family encompasses the gauges most commonly used in the literature, including the trivial, Euclidean, equitable, zero-sum, and wild-type gauges. For example, the zero-sum gauge is often used to characterize transcription factor binding specificities [19–21] and to compute contact maps from pairwise models of protein fitness landscapes [23, 24]. The wild-type gauge, by contrast, is standard when considering the impact of mutations relative to a specific reference sequence. See Posfai et al. [16] for details.

The hierarchical gauges, obtained by taking λ→ ∞, are a particularly useful subset of the λ, *π* gauges, one that includes both the zero-sum and wild-type gauges. In every hierarchical gauge, lower-order terms explain as much of the landscape’s variance as possible, while higher-order terms capture the residual variation. Whereas most λ, *π* gauges can be applied only to all-order models, hierarchical gauges can be applied to the larger class of hierarchical models as well. Moreover, the values of model parameters in these gauges have a natural interpretation: each parameter represents the average effect of introducing specific characters at specific positions, over and above the effect expected from lower-order terms (when sequences are drawn from *π*). One can therefore observe how a complex sequence-function relationship behaves in different regions of sequence space by choosing different *π*. For example, using a uniform distribution for *π* yields global averages, whereas concentrating *π* near specific sequences of interest can reveal local sequence requirements.

## Implementation

In this section, we formalize the gauge-fixing problem and describe the algorithms GaugeFixer uses to solve it.

### Generalized one-hot models

We consider sequences of length *L* constructed from an alphabet *A* = {*c*_1_, *c*_2_, …, *c*_*α*_} comprising *α* distinct characters, i.e., s ∈ *A* ^*L* 1^. Let *S* = {1, 2,…, *L*} denote the set of positions within *s*. Linear models of sequence-function relationships have the form

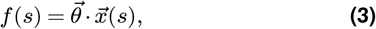

where 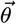 is a vector of model parameters and 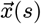 is a vector of sequence-dependent features.

We focus on one class of linear models, the generalized one-hot models described by Posfai et al. [16, 17]. In these models, each sequence-dependent feature is a binary indicator for a sub-sequence at a set of positions. Formally, each feature 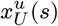 is specified by a subset of positions *U* ⊆ *S* and a subsequence *u* ∈ *A*^|*U* |^:

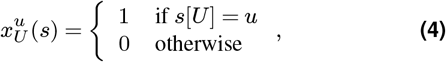

where *s*[*U*] indicates the subsequence of *s* that occurs at the positions *U*.^2^ We also require that 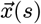 be equivariant under position-specific character permutations [17]. This means that if 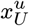 occurs in 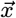 for some *U* and subsequence *u*, so must 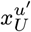 for every possible subsequence *u*′ ∈ *A*^|*U* |^. The set of features in a generalized one-hot model is therefore defined by an alphabet *A* and a collection of position sets *V* = {*U*_1_, *U*_2_,…, *U*_*n*_} via

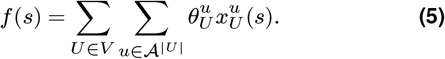

### All-order models and hierarchical models

All-order models are generalized one-hot models that include features for all possible subsequences at all sets of positions. This means that *V* includes every subset of positions *U* ⊆ *S*, i.e., *V* = *P* (*S*) is the powerset of *S*. We call *S* the generating set of *V*

An all-order model has *M* = (*α*+ 1)^*L*^ parameters. To see this, note that there are 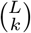 ways to choose a set of positions *U* of size *k*, and *α*^*k*^ possible subsequences u for each such set. Summing over *k* gives 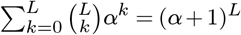 parameters. Alternatively, we can derive this parameter count by assigning to each feature 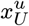 an index sequence of length *L* that has subsequence *u* at the positions in *U* and a wild-card character at all other positions. Because these index sequences are constructed from the alphabet, {∗ *c*_1_,…, *c*_*α*_}, which has size *α*+ 1, there are (*α*+ 1)^*L*^ possible index sequences, and hence the same number of parameters. This indexing approach underlies the tensor representation introduced by Posfai et al. [16] and used below.

Hierarchical models generalize all-order models. Each hierarchical model is defined by one or more generating sets *S*′ ⊆ *S*, whose subsets together make up *V*. Formally, if 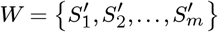 denotes the set of generating sets, then 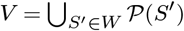 is the union of their powersets. Common examples of hierarchical models include additive models (generated by all position sets of size 1), pairwise models (all position sets of size 2), *K*-order models (all position sets of size *K*), nearest-neighbor models (all pairs of adjacent positions), and *K*-adjacent models (all windows of *K* adjacent positions).^3^

### Projection matrices for gauge-fixing

Posfai et al. [16] proposed the following approach for fixing the gauge of all-order models. Given an *M*-dimensional vector 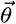 of initial parameter values, the gauge-fixed parameters 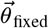 are obtained by multiplying 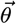 by an *M × M* projection matrix *P*:

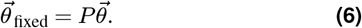

This matrix P is determined by λ and *π*. Specifically, it is constructed as a Kronecker product of position-specific matrices,

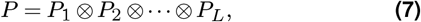

where P_*l*_ is the (α + 1) *×* (α + 1) matrix

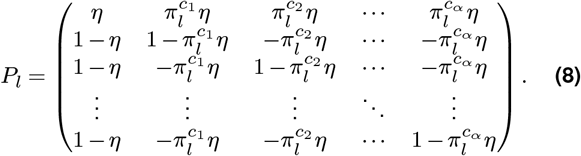

Here, the rows and columns of *P*_*l*_ are indexed by the characters ∗, *c*_1_, …, *c*_*α*_ (in that order) used to construct the index sequences, 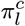 is the probability of character *c* occurring at position *l*, and η = λ/(1 +λ). See Posfai et al. [16] for details.

Hierarchical gauges, which correspond to *η* = 1, can be applied not just to all-order models but also to hierarchical models more generally. Conceptually, this is done by treating a hierarchical model as an all-order model on sequences of the same length, with unused parameters set to zero. Unlike other λ, *π* gauges, hierarchical gauges are guaranteed to leave these zeros in place, and therefore preserve the hierarchical form of the model [16]. As we show below, however, there is no need to construct the full all-order parameter vector to do this. Hierarchical models can therefore be gauge-fixed even when *L* is far too large for all-order models to be practical.

### Gauge-fixing algorithm for all-order models

The Kronecker product form of *P* in Eq. 7 allows P 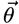 to be computed without explicitly constructing *P* [14]. To do this, we first reshape 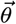 into an *L*-dimensional tensor having size *α* + 1 along each dimension. The expression in Eq. 6 then becomes

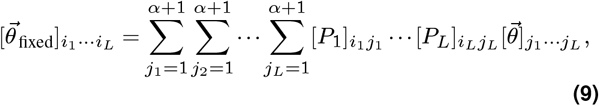

where each index i_*l*_ runs from 1 to *α* + 1. We evaluate Eq. 9 iteratively: first we define 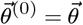; then for each l = 1, 2,…, *L* we compute

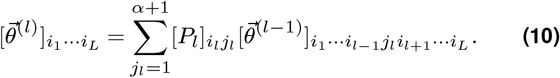

Iterating this expression yields 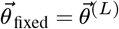. This computation is formalized in Algorithm 1. Note that this algorithm uses an intermediate parameter vector 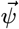 to avoid storing all 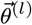 simultaneously.

#### Algorithm 1

Fixing the gauge of all-order models

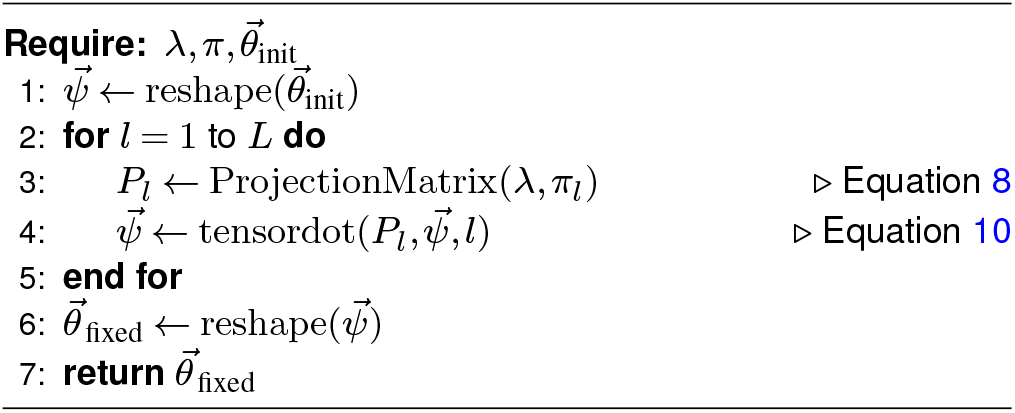

We now derive the computational complexity of Algorithm 1. For each choice of *l* and indices *i*_1_, …, *i*_*L*_, it takes *α*+1 multiplication-addition operations to compute the right-hand side of Eq. 10. As there are *L* values of *l* and (*α* + 1)^*L*^ possible combinations of indices, the algorithm has computational complexity *O*(*L*(*α* + log(α+1) 1)^*L*+1^). In terms of *M* and *α*, this is 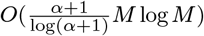, which for fixed *α* becomes *O*(*M* log *M*). By contrast, standard matrix multiplication by *P* requires *O*(*M* ^2^) operations.

The memory requirements of Algorithm 1 are also dominated by line 4, which implements Eq. 10. For each choice of *l*, this step requires storing 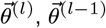, and *P*_*l*_, which together amount to 2*M* + (*α* + 1)^2^ numbers. Since *M* ≥ (*α* + 1)^2^ whenever *L* ≥ 2, and the algorithm never stores more than two *M*-sized vectors at once, these requirements scale as *O*(*M*). By contrast, storing *P* itself requires storing *O*(*M* ^2^) numbers in memory.

### Gauge-fixing algorithm for hierarchical models

To express the parameters of a hierarchical model in a hierarchical gauge, we use the fact that these parameters can be decomposed into a sum of the parameters of multiple all-order models, each restricted to one of the generating position sets. Let 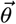 be the parameter vector of the hierarchical model expressed as an (*α* + 1)^*L*^-dimensional all-order parameter vector (with unused parameters set to zero), and let W denote the set of generating sets used to define it. We first decompose 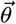 as

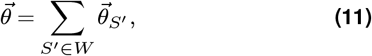

where each 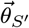 is an all-order parameter vector for the position set *S*^*′*^, i.e., has nonzero parameters 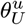 only for *U* ⊆ *S*^*′*^. The gauge-fixed parameters are then computed as

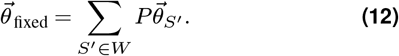

Note that there are often many valid ways to carry out this decomposition due to the generating sets in *W* overlapping. Because *P* is a linear operator, however, the result is independent of which decomposition is used. Each term on the right-hand side can be computed using Algorithm 1 with only the Kronecker factors *P*_*l*_ for positions *l* ∈ *S*^*′*^. Thus, there is never any need to represent the full (*α* + 1)-dimensional vectors 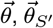, or 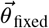.

Algorithm 2 formalizes this procedure. Here, 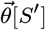 indicates an (*α*+ 1)^|*S*^*′*^|^-dimensional vector composed only of the elements 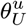 for which U ⊆ *S*′, and *π*[*S*′] denotes the marginal distribution over subsequences on *S*′. By zeroing out 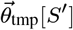 at line 7, the algorithm selects one particular valid decomposition: each parameter 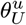 is assigned to the first generating set *S*′ for which U ⊆ *S*′. This iterative form also avoids the need to store the nonzero elements of all 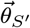 simultaneously.

#### Algorithm 2

Fixing the gauge of hierarchical models.

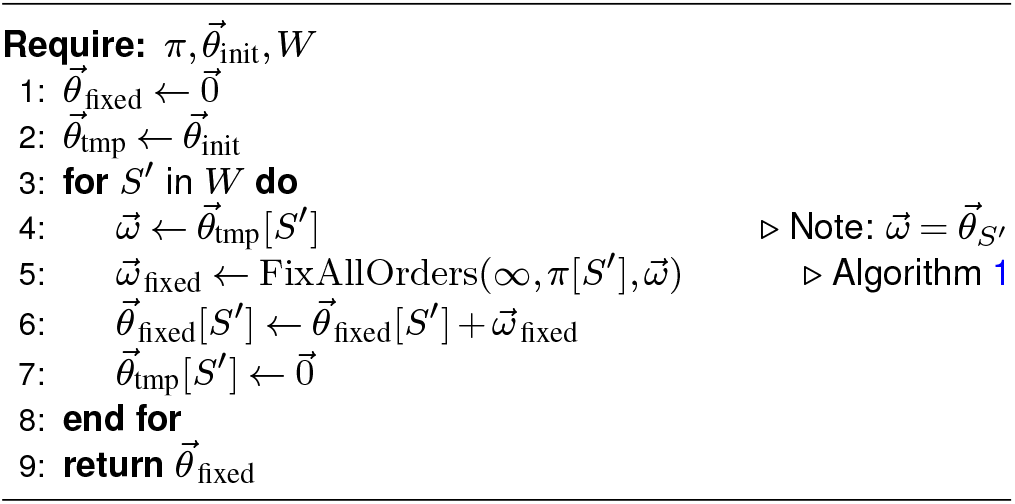

The memory requirements of this algorithm scale as *O*(*M*), just as for Algorithm 1. The computational requirements, however, depend on the size of *W* and on the size of each *S*′ ∈ *W*, both of which are model-specific. For *K*-order models specifically, the generating sets consist of all possible subsets of *K* positions; this means that 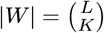 and |S | = *K* for all *S* ∈ *W*. The primary computational expense occurs in line 5, which is carried 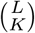 out times. Each execution requires *K*(*α* + 1)_*K* + 1_ operations, following the analysis above. By comparison, the number of parameters is 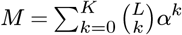. Assuming *K* ≪ *L*, we can approximate 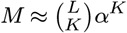. Holding *α* and *K* fixed while increasing *L* thus causes the computational complexity to scale as *O*(*M*). *K*-adjacent models behave similarly.

## Results

### Performance benchmarking

We benchmarked GaugeFixer’s implementations of Algorithms 1 and 2 against standard matrix multiplication using full-sized projection matrices. We did this for all-order and pairwise models of varying size. As expected, GaugeFixer achieved orders-of-magnitude improvements in both runtime and memory usage (Figure 2). In particular, standard matrix multiplication became impractical for models having more than about ten thousand parameters, whereas GaugeFixer processed models having millions of parameters in just a few seconds.

**Fig. 2.**
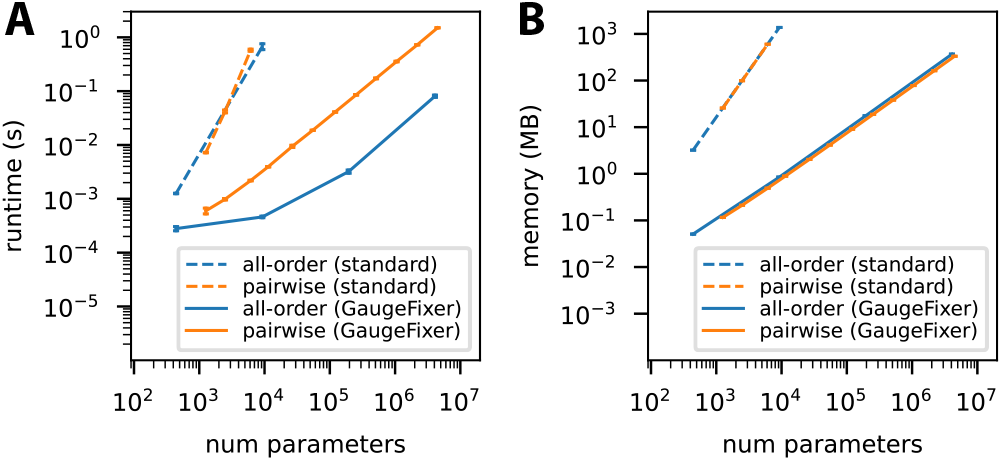
Benchmarking results. Shown are the runtime (A) and peak memory (B) requirements for GaugeFixer and for standard matrix multiplication, plotted as a function of the number of model parameters. Tests were carried out for all-order models (*L* = 2 to 5) and pairwise models (*L* = 3 to 150) on protein sequences (*α* = 20). Each data point represents the mean and standard deviation over 10 gauge-fixing computations performed on a standard laptop computer. No execution-time limit was imposed, but benchmarking computations for standard matrix multiplication were restricted to models whose projection matrix required less than 2 GB of memory.

### Application to the Shine-Dalgarno fitness landscape

To illustrate the utility of GaugeFixer, we analyzed an empirical fitness landscape for translation initiation. Recognition of the translation start site in many bacteria is mediated by the Shine-Dalgarno (SD) sequence, a motif that occurs in the 5’ UTR of mRNA and functions by base-pairing with the 3’ tail of the 16S ribosomal RNA [42]. To measure this sequence-function relationship, Kuo et al. [40] randomized a 9-nt region within the 5’ UTR of the *dmsC* gene, spanning positions − 13 to − 5 relative to the start codon. They used these variant UTRs to drive translation of a fluorescent protein in *Escherichia coli*, then quantified expression using fluorescence-activated cell sorting followed by deep sequencing. This yielded activity measurements for 257,565 of the 4^9^ = 262,144 possible 9-nt sequences.

Recently, Martí-Gómez et al. [14] extended this landscape to all possible 9-nt sequences using variance component regression, a Bayesian inference method based on Gaussian processes [11]. This method estimates the variance attributable to genetic interactions of each order, then uses these estimates to define a prior over landscapes. Having inferred a combinatorially complete landscape, they applied the visualization method of McCandlish [43]. Doing so, they observed that the SD landscape contains multiple prominent fitness peaks corresponding to the canonical AGGAG motif appearing in different registers (i.e., starting at different positions upstream of the start codon).

Here we used GaugeFixer to characterize the landscape around each fitness peak. We represented the combinatorially complete landscape of Martí-Gómez et al. [14] as an all-order model having (4 + 1)^9^ = 1,953,125 parameters expressed in the trivial gauge (in which only the 4^9^ parameters of order 9 are nonzero). Then for each register *i*,^4^ we defined a distribution π that fixes the core motif *u* = AGGAG at that register while randomizing the other positions within the 9-nt variable region (Figure 3A).^5^ We next used GaugeFixer to express these parameters in the hierarchical gauge corresponding to each choice of π. These gauges are examples of the generalized wild-type gauges described by Posfai et al. [16], in which a reference sequence is specified at only a subset of positions. Under such a gauge, the parameters describe the average effects of mutations away from the reference sequence, computed over random backgrounds at the remaining positions. We emphasize that, although the resulting parameter values differ across gauges, they are alternative representations of the same underlying model and therefore yield identical predictions.

**Fig. 3.**
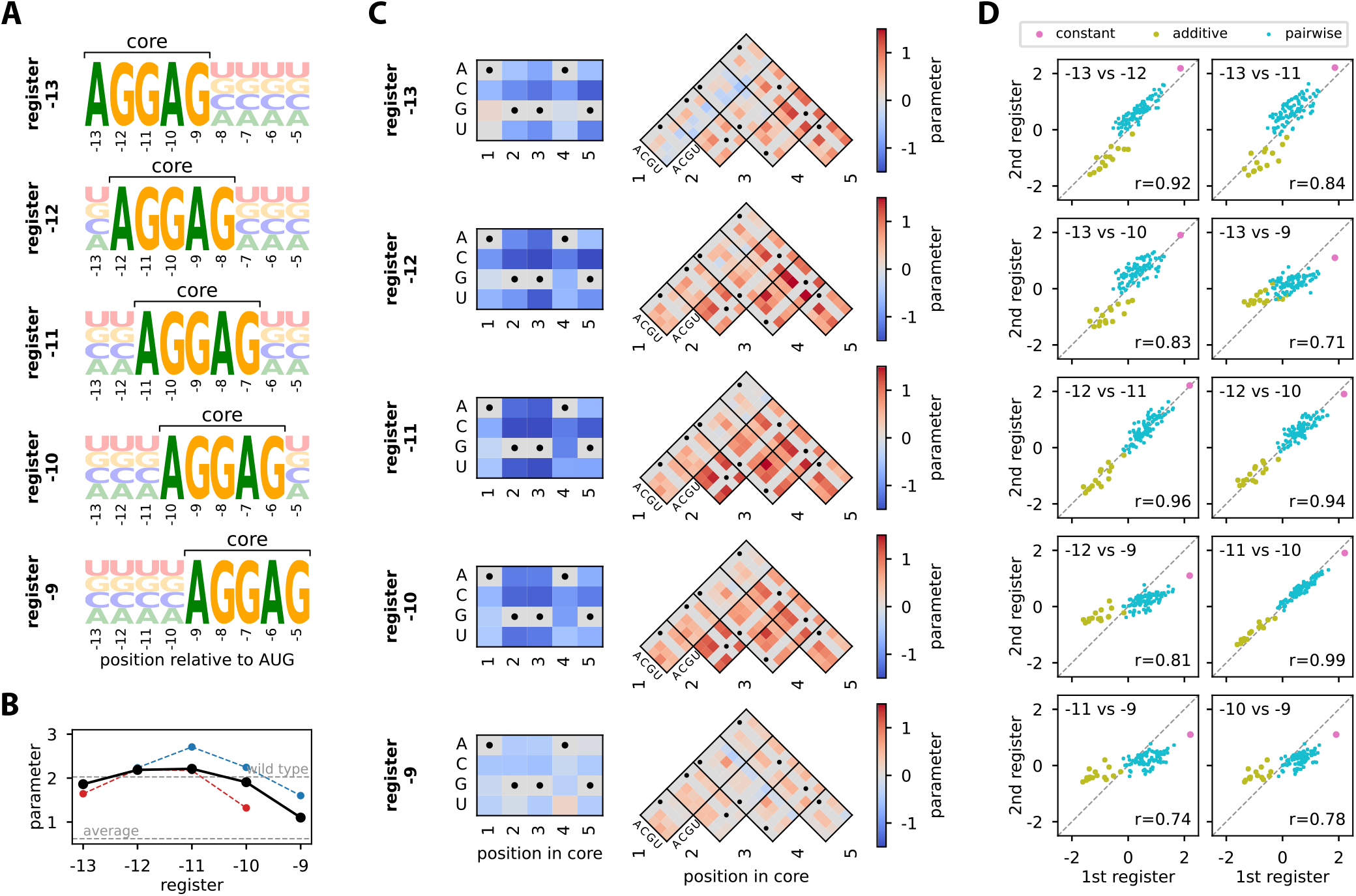
Application of GaugeFixer to a fitness landscape for translation initiation. (A) Sequence logos [41] showing the probability distributions *π* corresponding to different peaks in the Shine-Dalgarno landscape measured by Kuo et al. [40] and modeled by Martí-Gómez et al. [14]. Each distribution has a fixed AGGAG motif at a specific register, i.e., starting position relative to the start codon. (B,C) Model parameters expressed in hierarchical gauges corresponding to different choices of *π*. (B) Values of the constant parameters corresponding to the *π* distributions illustrated in panel A (black). Also shown are the values resulting from *π* distributions with extended core motifs: GAGGAG (blue) and AGGAGU (red). Both extended-motif series are plotted against the register of the AGGAG core motif. Horizontal dashed lines represent the average activity across all possible sequences (lower) and the activity of the wild-type sequence AAGGAGGUG (upper). (C) Gauge-fixed additive (left) and pairwise (right) parameters in the same gauges. Black dots mark the reference nucleotide at each position of the core motif, as indicated in panel A. (D) Scatterplots comparing constant, additive, and pairwise parameter values obtained for the five different registers in panel A.

The constant parameter represents the mean activity of sequences with AGGAG in the specified register. We found that this value was highest for registers − 12 and − 11 and markedly reduced for register − 9 (Figure 3B). This pattern is consistent with the known spacing requirements for translation initiation: register − 12 is the established optimum [44], whereas register − 9 places the SD motif closer to the start codon than occurs naturally in nearly all bacterial genes [45].

The dependence of the constant parameter on register could reflect the intrinsic positional preference of the core motif, the influence of the fixed sequence flanking the variable region, or both. Every assayed sequence carries a fixed G at position − 14 and a fixed U at position − 4, just outside the variable region. The effects of these characters cannot be read directly from the model. They can, however, be probed indirectly: extending the core motif to GAGGAG or AGGAGU places a G or a U immediately adjacent to the core, and the resulting constant parameter reports the mean activity of sequences carrying that character at that position. We therefore computed constant parameters under hierarchical gauges defined by these extended motifs at every register where they fit within the variable region (Figure 3B). Neither extended motif can be placed at the register where its added character would coincide with the corresponding fixed position in the assay: AGGAGU does not fit at register − 9, nor GAGGAG at register − 13. The neighboring registers, − 10 and − 12 respectively, therefore provide the closest available proxies. At register − 10, the mean activity of sequences containing AGGAGU was substantially lower than that of sequences containing AGGAG alone; a smaller decrease was observed at register − 13, and essentially none at registers − 12 and − 11. This suggests that the low average activity of sequences carrying the core motif at register − 9 is partly attributable to the negative contribution of the fixed − 4U, although this contribution is itself register-dependent. By contrast, the extended motif GAGGAG raised mean activity at registers − 11 through − 9, but not at register − 12. The fixed − 14G is therefore unlikely to explain why the constant parameter is lower at register − 13 than at registers − 12 and − 11; an additional upstream G appears to be beneficial only when the core motif lies closer to the start codon.

The additive parameters, shown in Figure 3C (left), reveal how individual nucleotide mutations away from the AGGAG core motif affect activity in each register. As expected, these mutations were overwhelmingly deleterious. The pattern of these effects was also remarkably consistent across registers. Similarly, the pairwise interaction parameters (Figure 3C, right) capture the effects of mutating pairs of nucleotides within the core motif beyond what additive effects alone predict. These interactions also varied little across registers and were predominantly positive, indicating that combinations of mutations tend to be less deleterious than expected from their individual effects. Such positive interactions are a hallmark of global epistasis [1, 9], and are consistent with a previously proposed biophysical model for Shine-Dalgarno sequence activity [14]. Both the additive and the pairwise parameters were, however, smaller in magnitude at the extreme registers − 13 and − 9, indicating that mutations to the core motif have less effect on activity when the motif is poorly positioned.

Finally, we compared the constant, additive, and pairwise parameters in the core motif across registers (Figure 3D). Neighboring registers tended to have similar parameters, while distant registers diverged more, suggesting that the sequence preferences of the ribosome change gradually with register.

## Discussion

GaugeFixer provides an efficient tool for fixing the gauge of commonly used quantitative models of sequence-function relationships. By leveraging the Kronecker product form of projection matrices, GaugeFixer dramatically reduces both the time and the memory required for this computation. This makes it possible to carry out, in seconds, gauge-fixing computations that would otherwise be impractical. We demonstrated this capability on an empirical landscape for translation initiation described by a model with nearly two million parameters. The results illuminated how the sequence requirements of the Shine-Dalgarno motif depend on its distance from a gene’s start codon, revealing remarkable consistency but also fine-scale variation across registers.

We wish to emphasize that gauge fixing is fundamentally different from parameter inference. Both procedures are essential for modeling sequence-function relationships, but they serve distinct purposes. Inference identifies parameters that make a model’s predictions best fit one’s data without regard to how those parameters are interpreted. Gauge fixing, by contrast, alters model parameters in ways that have no effect on model predictions but are important for their interpretation. GaugeFixer respects this distinction by operating only on precomputed parameter vectors; it is not a tool for inferring parameter values from data.

A related but more subtle point is that inference procedures using certain regularization schemes (or prior distributions in Bayesian contexts) can themselves yield parameters in specific gauges [16, 46]. This does not, however, eliminate the need for post hoc gauge fixing if model parameters are to be meaningfully interpreted. Standard L2 regularization, for example, yields parameters in the Euclidean gauge, whereas researchers often wish to interpret parameters in the zero-sum or wild-type gauges.

Gauge fixing is conceptually related to neural network interpretability methods such as “global importance analysis” [47] and “in silico marginalization” [48, 49]. These methods involve evaluating a model of interest on a large number of sequences, parts of which are held fixed and parts of which are randomized. While these interpretability methods can be applied to any model, they require substantial computation and yield only stochastic estimates of the effect of a sequence pattern averaged over random backgrounds. Gauge fixing, by contrast, provides exact results not only for the average effect of a sequence pattern in random backgrounds (via the specification of *π*), but also for the expected effects of mutations and their combinations.

Although GaugeFixer operates only on specific classes of linear models (all-order and hierarchical models), it can still be used to characterize nonparametric or nonlinear models (e.g., Gaussian process models or deep neural networks). To do this, one first evaluates the model of interest on every possible sequence. The resulting landscape is then represented as an all-order model, and GaugeFixer is applied to the corresponding parameters. This is what we did for the Shine-Dalgarno landscape analyzed above (Figure 3). Because this approach requires explicitly representing the combinatorially complete landscape, it can be used only when sequences are short enough for that to be computationally feasible. We note that recent theoretical results, however, suggest a way of overcoming this limitation for certain Gaussian process models [46].

Finally, Algorithms 1 and 2 can also be used for purposes other than gauge fixing. In particular, the Walsh-Hadamard transformation of sequence-function landscapes [33, 50] and its generalizations [20, 39, 51, 52] involve multiplying an *α*^*L*^-dimensional vector (representing a combinatorially complete landscape) by a dense matrix constructed as a Kronecker product of position-specific matrices. Algorithm 1 allows these transformations to be performed far more efficiently than by standard matrix multiplication. Algorithm 2, moreover, extends these transformations to landscapes that are defined by hierarchical models (e.g., K-order or K-adjacent models), rather than by combinatorially complete sets of measurements. This latter capability should allow these methods to be applied to sequences that are much longer than is computationally feasible using published approaches. We have incorporated these other landscape-analysis approaches into GaugeFixer alongside the package’s core gauge-fixing functionality.

## Conclusions

GaugeFixer provides a high-performance software library for fixing the gauge of commonly used sequence-function models. This fills an important gap in the computational tools available for interpreting complex sequence-function relationships, including landscapes with high-order genetic interactions or multiple fitness peaks.

## Availability and requirements

**Project name:** GaugeFixer

**Project home page:** https://github.com/jbkinney/gaugefixer

**Documentation:** https://gaugefixer.readthedocs.io

**Operating system(s):** Platform independent **Programming language:** Python

**Other requirements:** Python ≥ 3.10; installable via the pip package manager

**License:** MIT

**Any restrictions to use by non-academics:** None

## List of Abbreviations

5’ UTR: 5’ untranslated region
DNA: deoxyribonucleic acid
RNA: ribonucleic acid
SD: Shine-Dalgarno

## Declarations

### Ethics approval and consent to participate

Not applicable.

### Consent for publication

Not applicable.

### Availability of data and materials

GaugeFixer source code is freely available under an MIT license at https://github.com/jbkinney/gaugefixer. Documentation is hosted at https://gaugefixer.readthedocs.io. Code and data for reproducing the analyses in this manuscript are available at https://github.com/jbkinney/25_gaugefixer. The Shine-Dalgarno fitness landscape analyzed here was originally measured by Kuo et al. [40] and modeled as described in Martí-Gómez et al. [14].

### Authors’ contributions

C.M-G. and J.B.K. conceived the project, wrote the package, and performed the research. C.M-G., D.M.M., and J.B.K. wrote the manuscript. D.M.M. and J.B.K. supervised the research. All authors read and approved the final manuscript.

## Acknowledgements

Not applicable.

## Footnotes

1 GaugeFixer also supports sequences built using different alphabets at different positions, with no need for these alphabets to be of the same size. The formulas in this section generalize in a straightforward way to accommodate this more general use case.

2 Sets of positions are called “orbits” in Refs. [16, 17] due to group-theoretic considerations. Such considerations do not concern us here, so we have opted to use more transparent terminology.

3 Posfai et al. [17] adopted a different definition of *K*-order and *K*-adjacent models, one that makes these models non-hierarchical. We favor the present definition, which is hierarchical and thus amenable to gauge fixing using hierarchical gauges.

4 Throughout this section we index sequence positions *l* by their offset from the start codon, so that *l* runs from − 13 to −5 rather than from 1 to 9. We also denote by *i* the register of the AGGAG core motif, so that I runs from −13 to −9.

5 Formally, 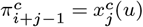 for 1 ≤ *j* ≤ 5, 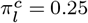 for *l* < *i* or *l* > *i*+ 4.

